## Supplementary figures and images for "Mgl2^+^ cDC2s coordinate fungal allergic airway type 2, but not type 17, inflammation"

### Supplementary Figure 1

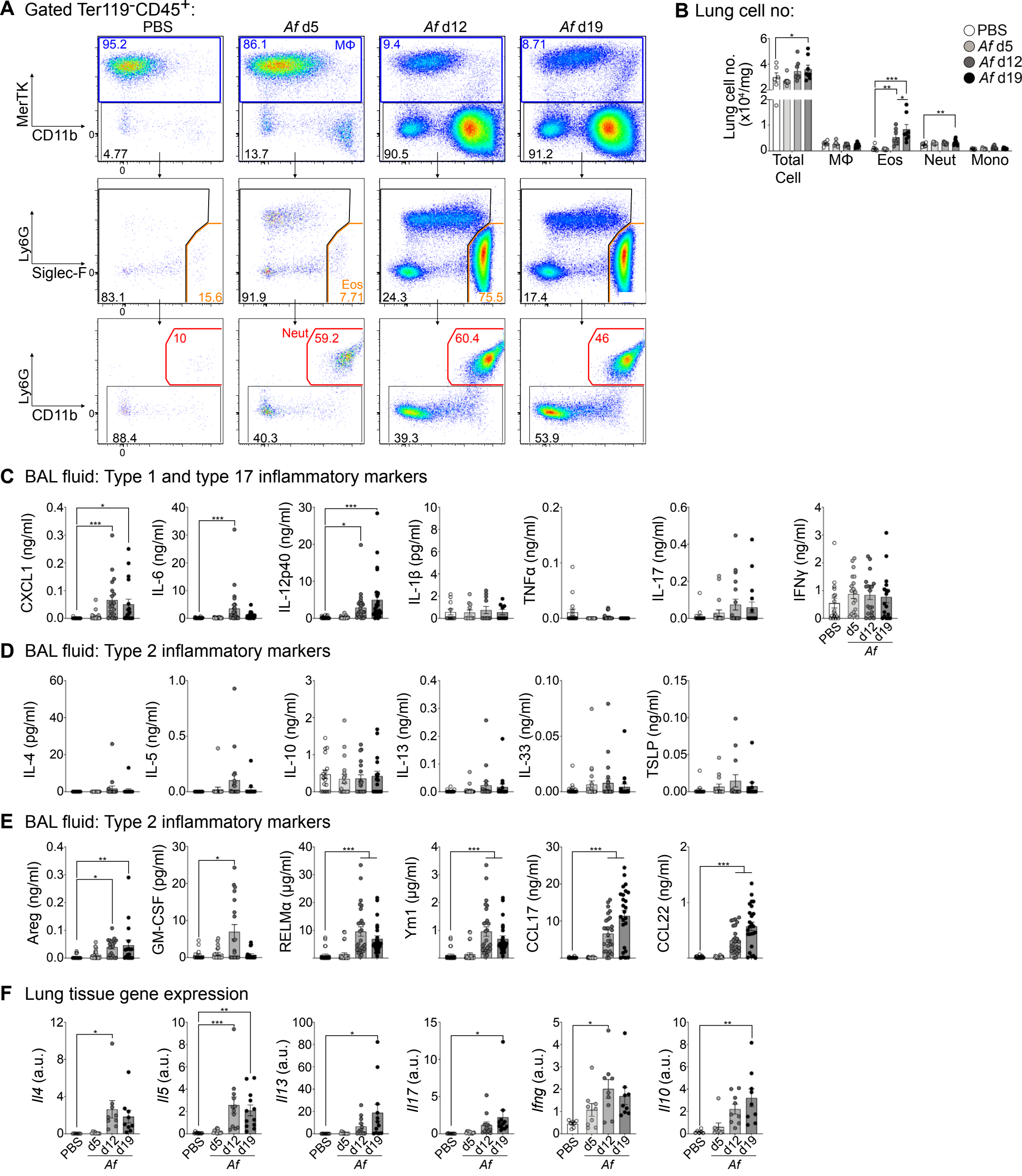


**Supplementary Figure 1**

### Supplementary Figure 2

**Lymphoid BAL cell gating**

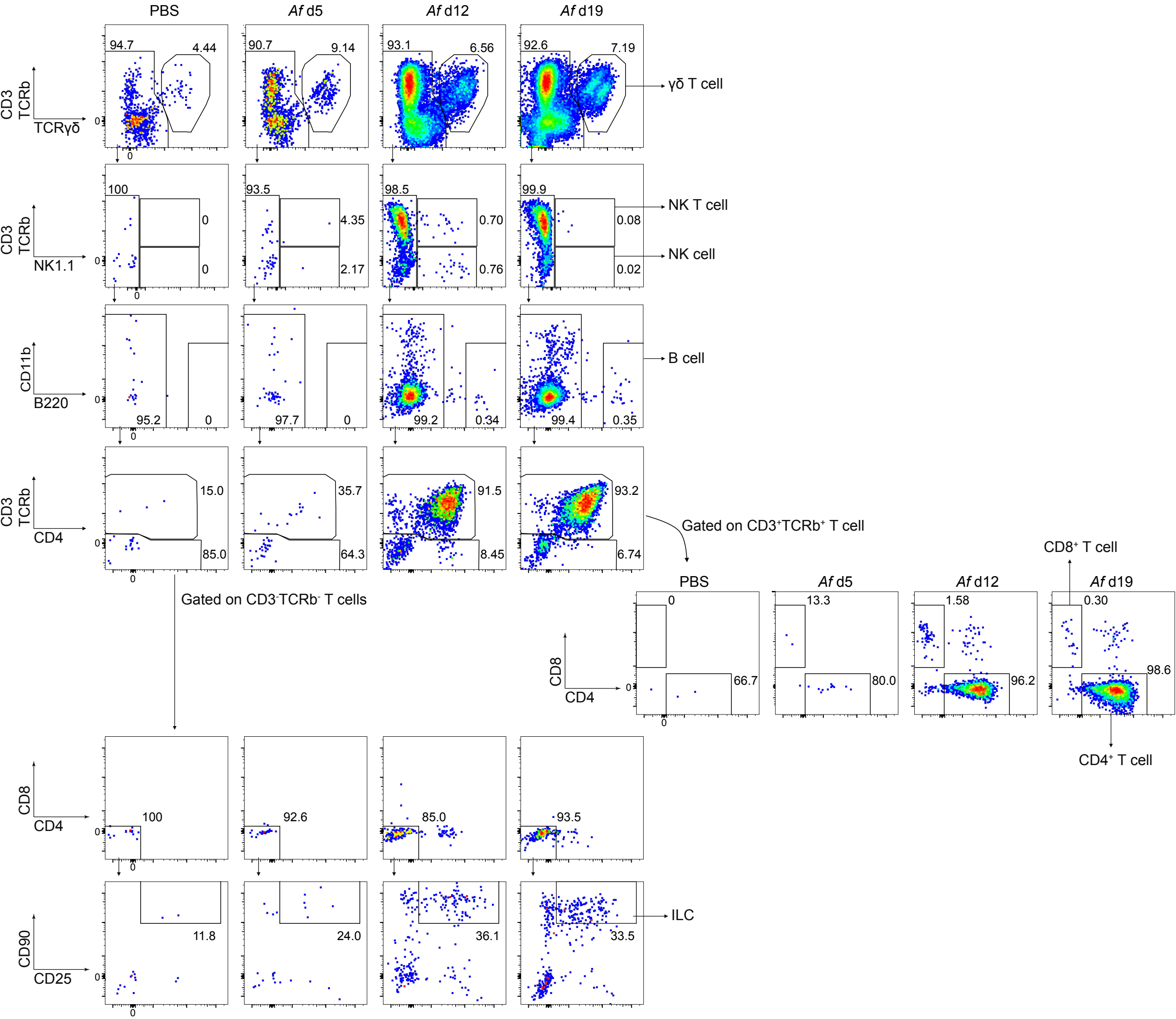


Supplementary Figure 2

### Supplementary Figure 3

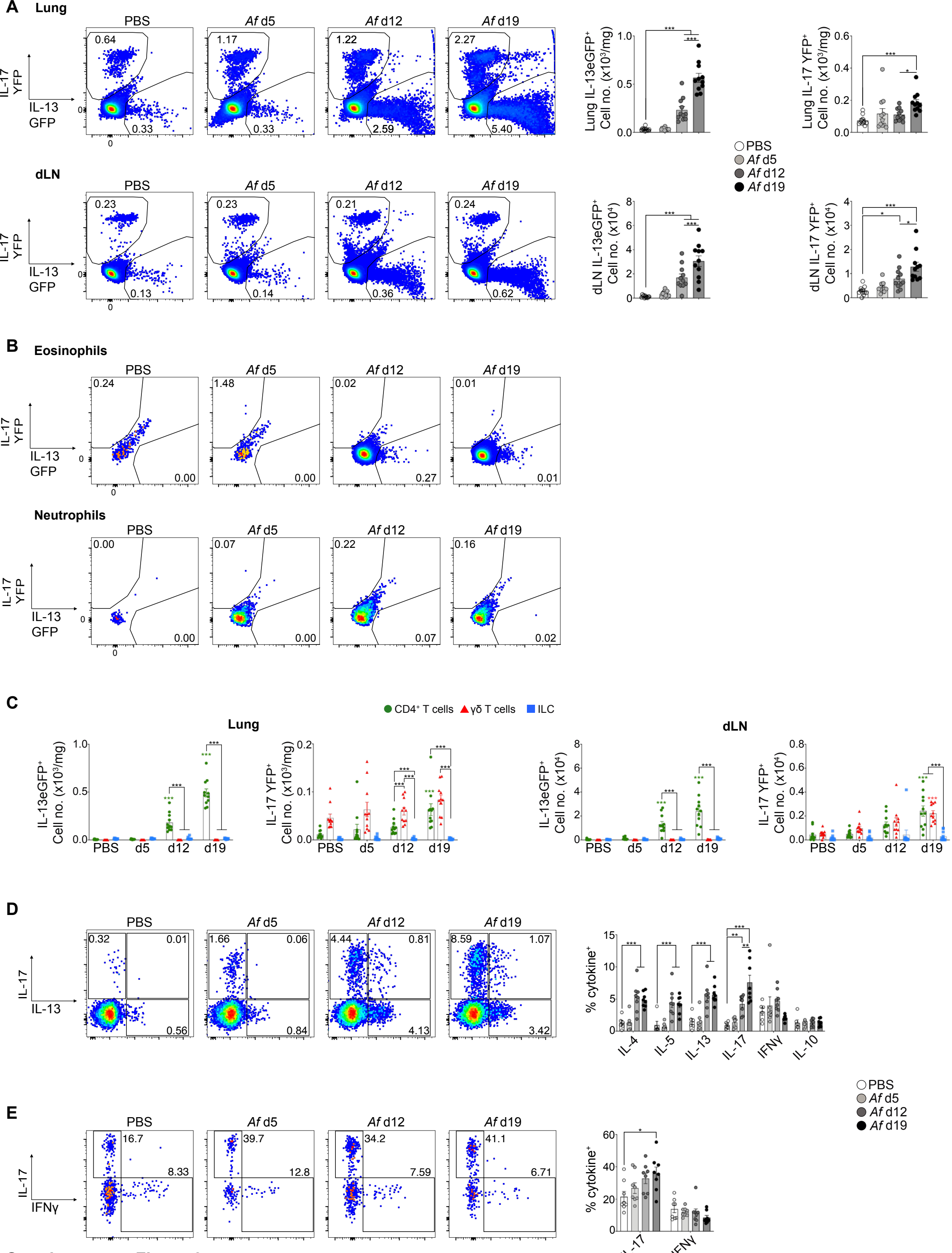


Supplementary Figure 3

### Supplementary Figure 4

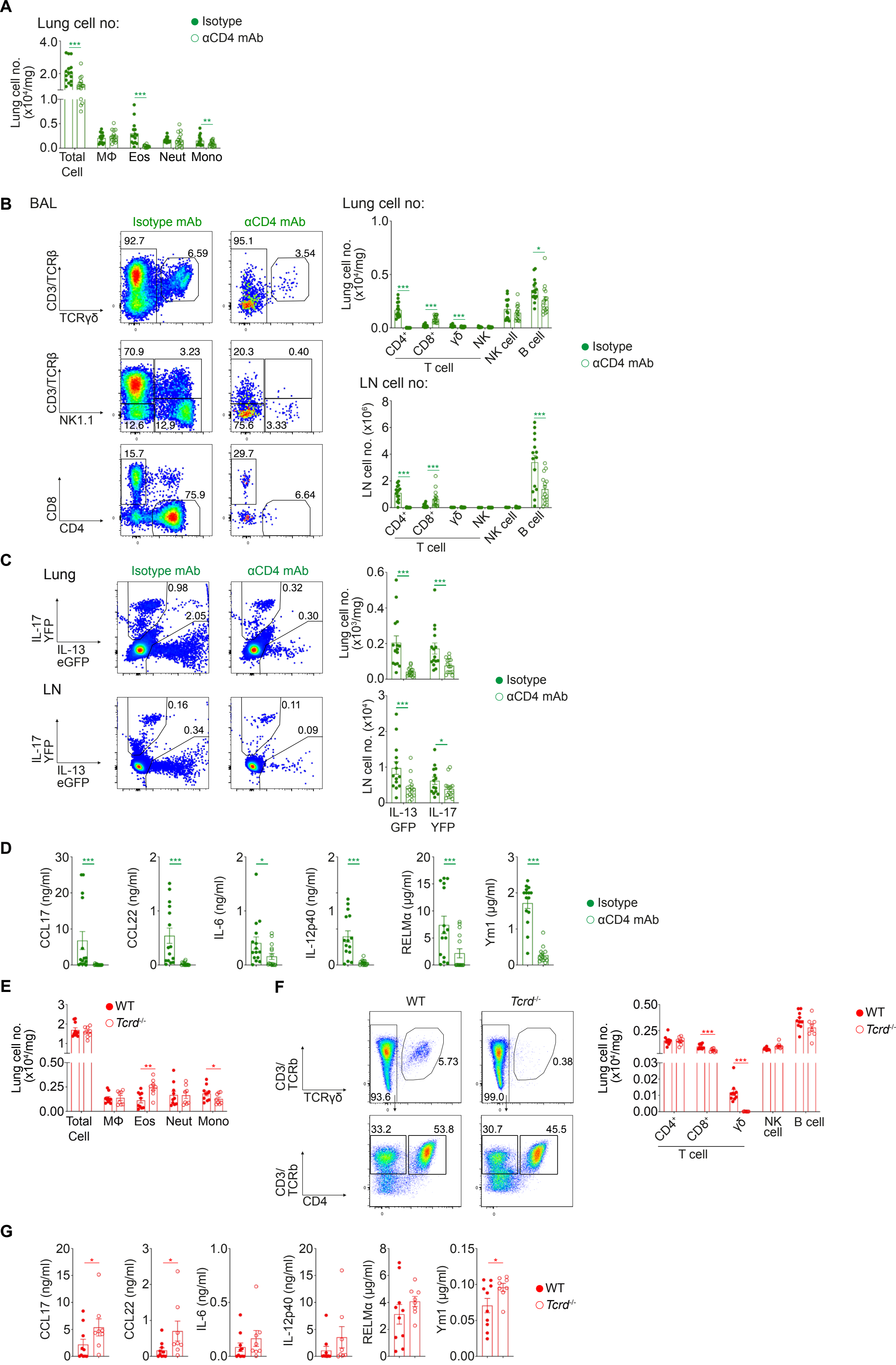


Supplementary Figure 4

### Supplementary Figure 5

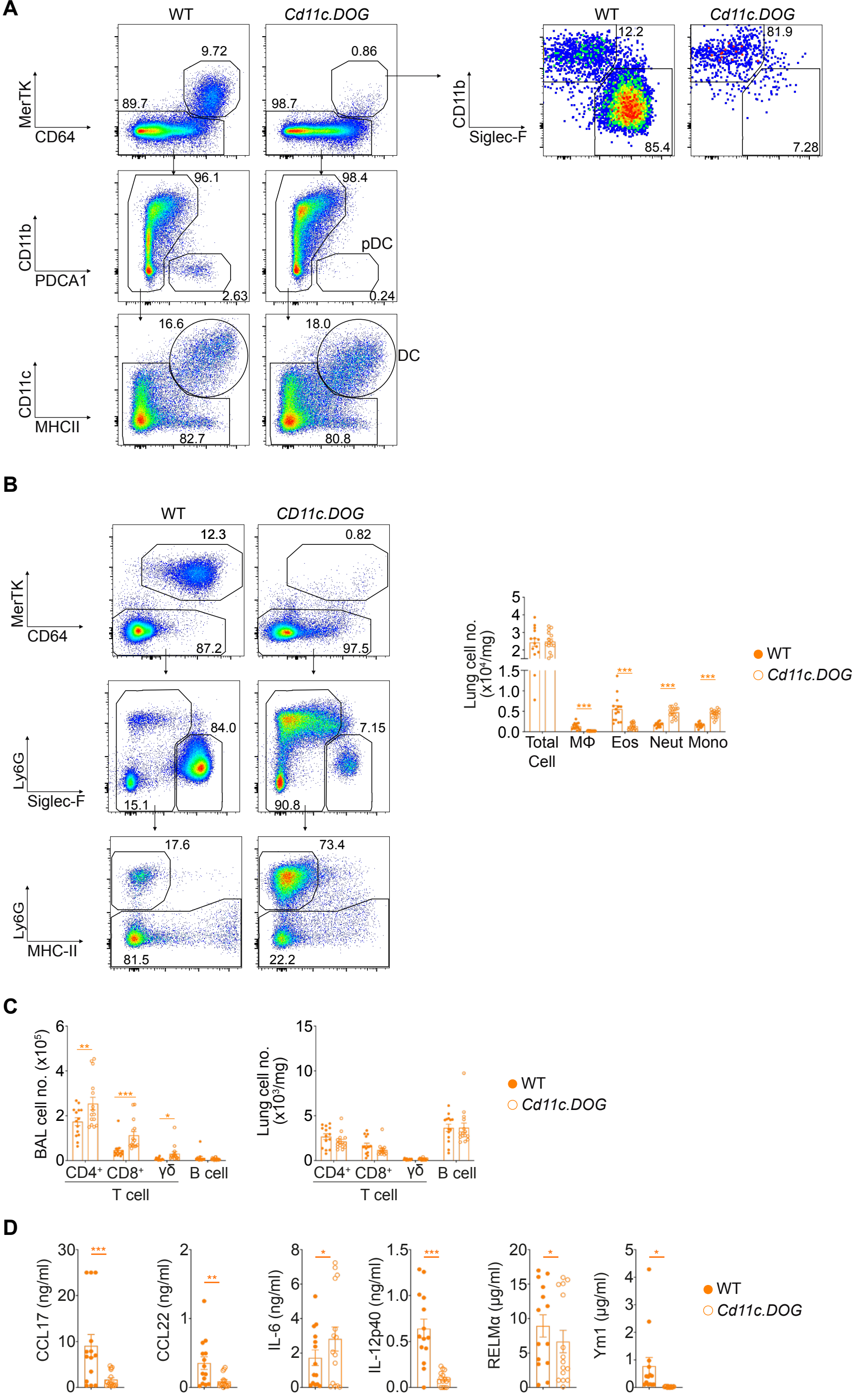


**Supplementary Figure 5.**

### Supplementary Figure 6

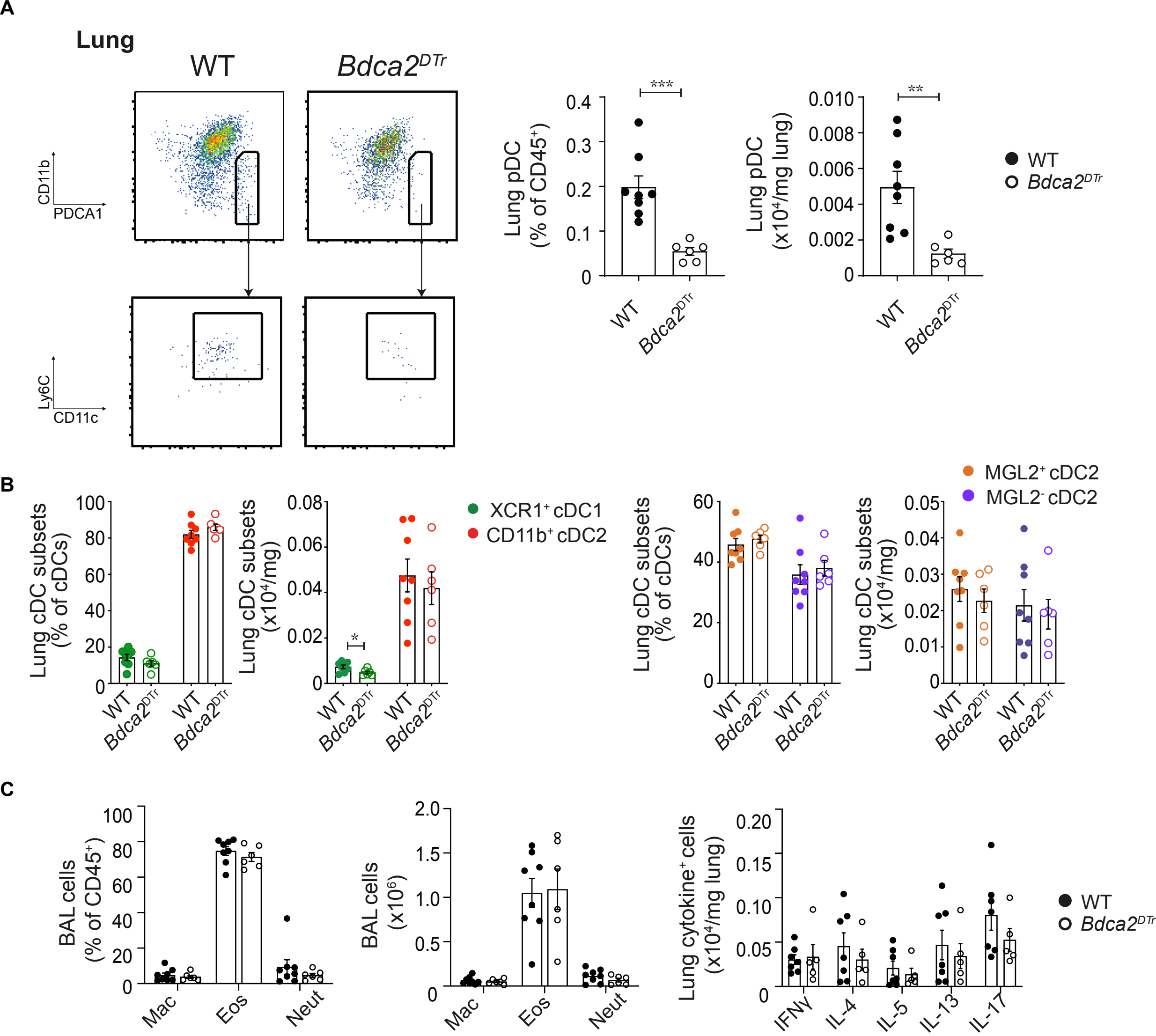


**Supplementary Figure 6.**

### Supplementary Figure 7

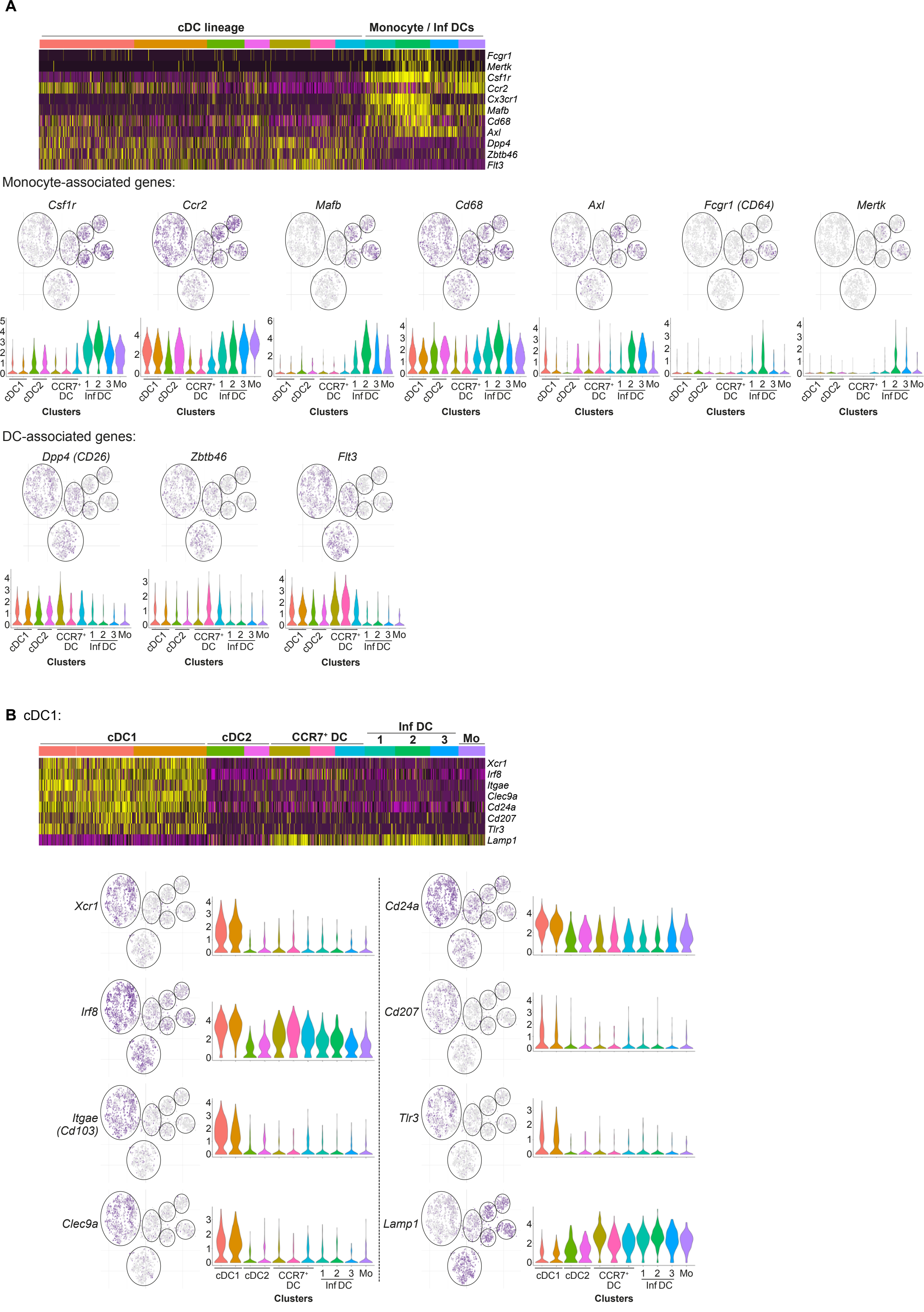


Supplementary Figure 7

### Supplementary Figure 8

**A** cDC2:

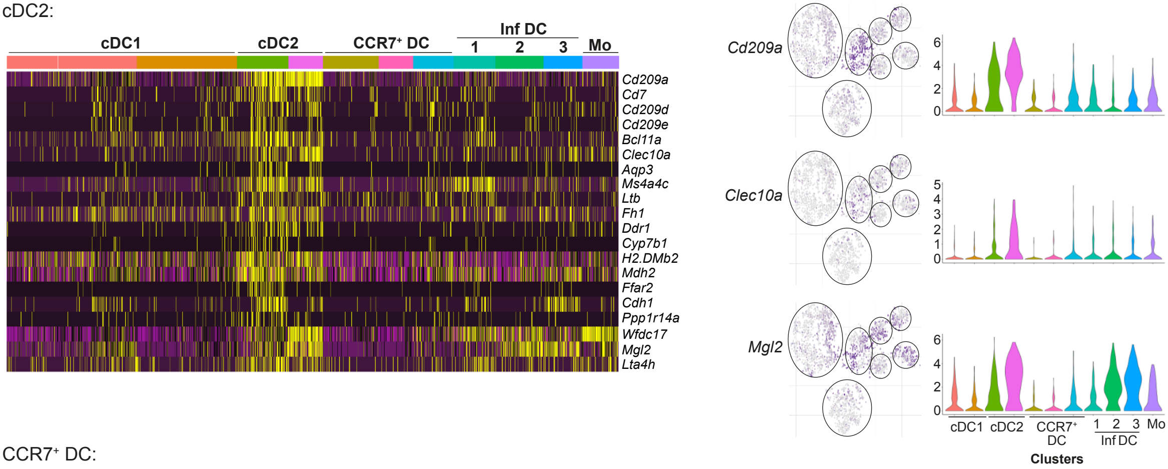


**B** CCR7⁺ DC:

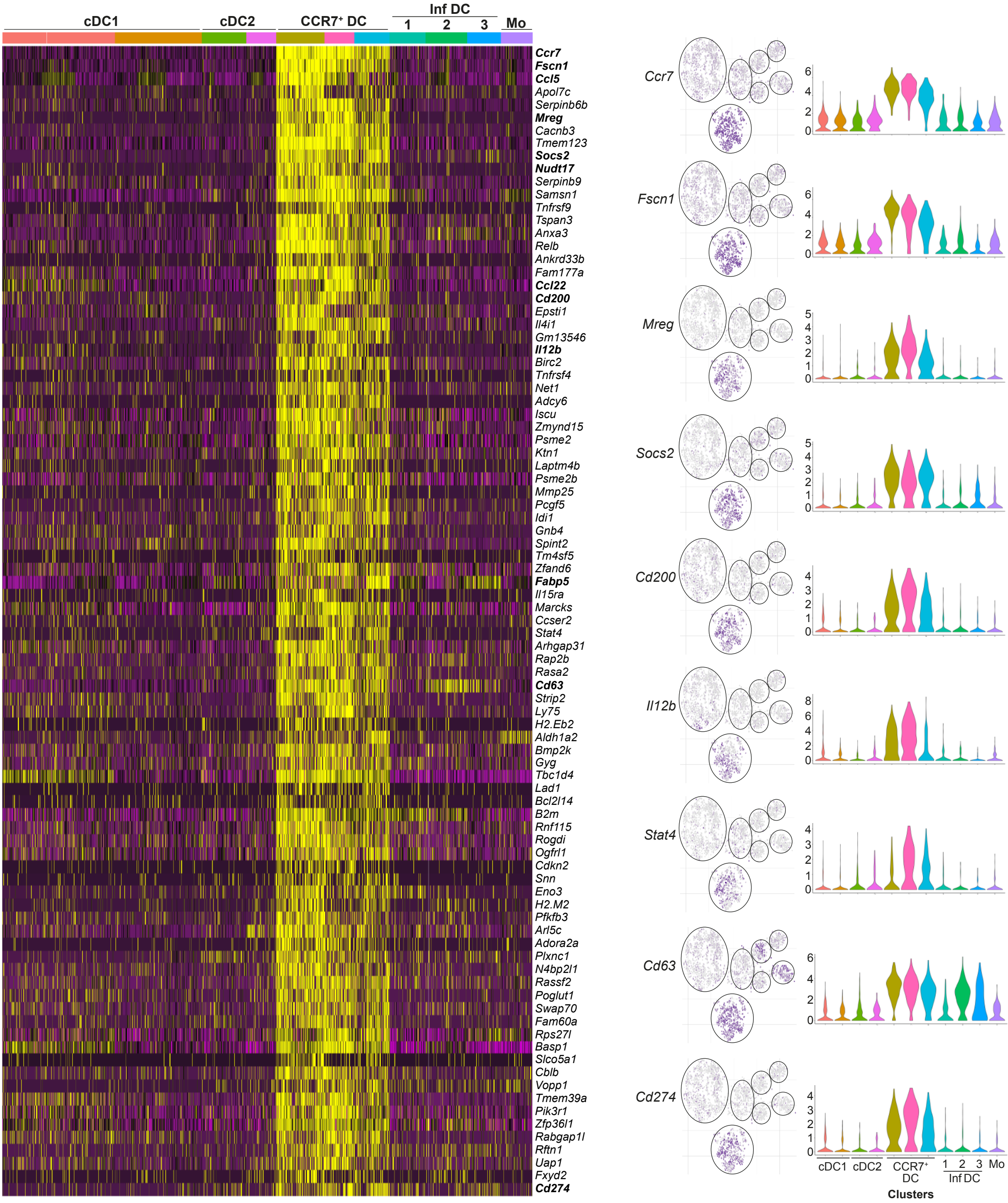


**Supplementary Figure 8**

### Supplementary Figure 9

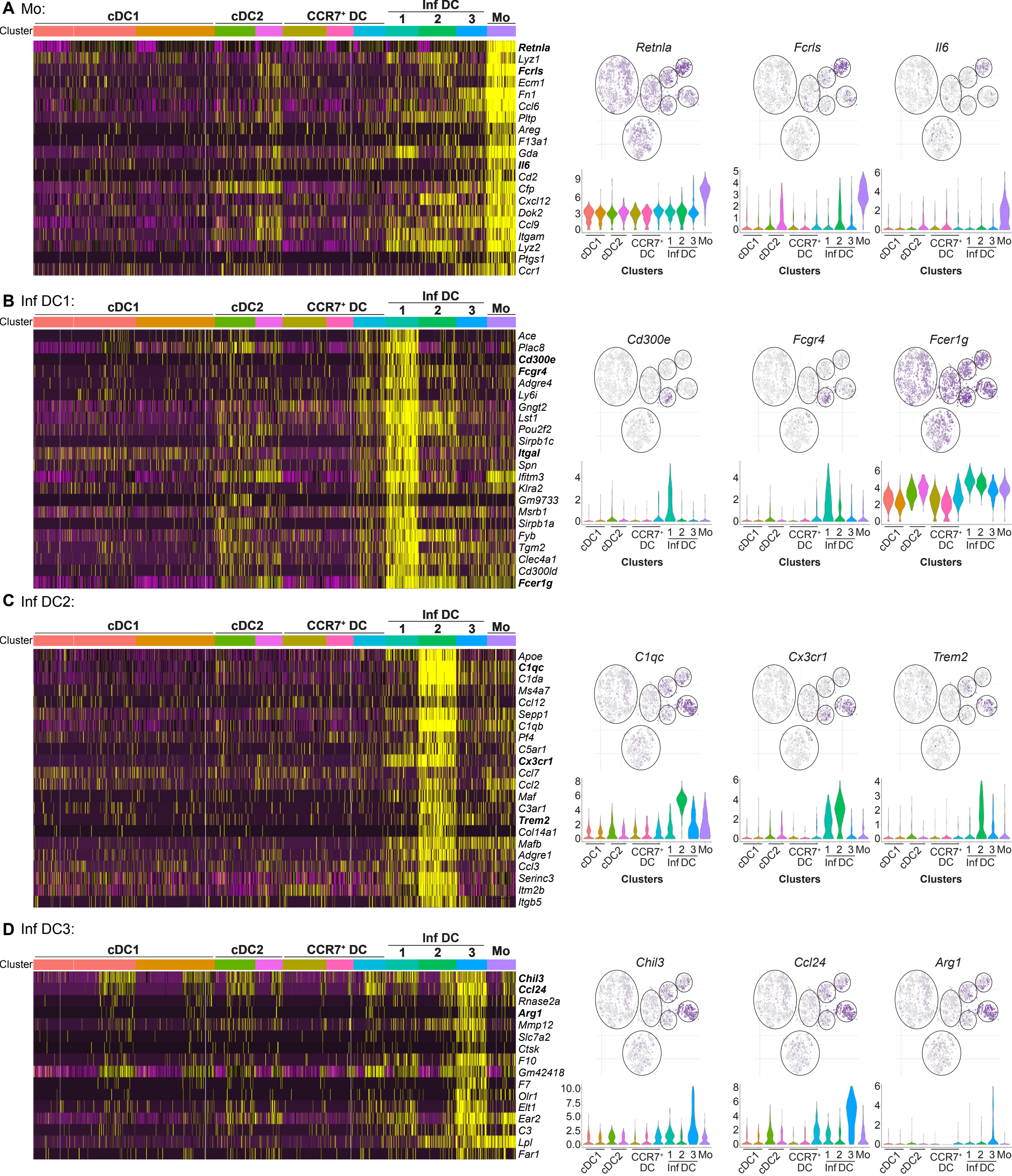


**Supplementary Figure 9**

### Supplementary Figure 10

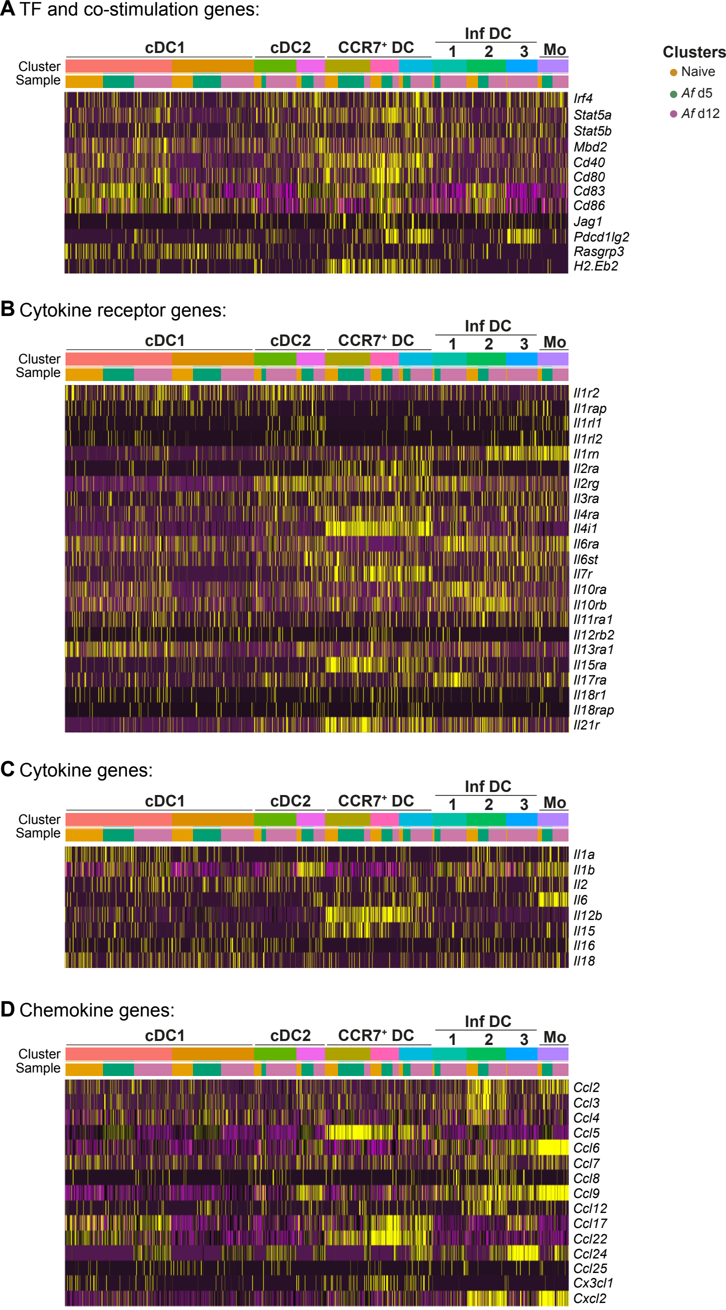


**Supplementary Figure 10**

### Supplementary Figure 11

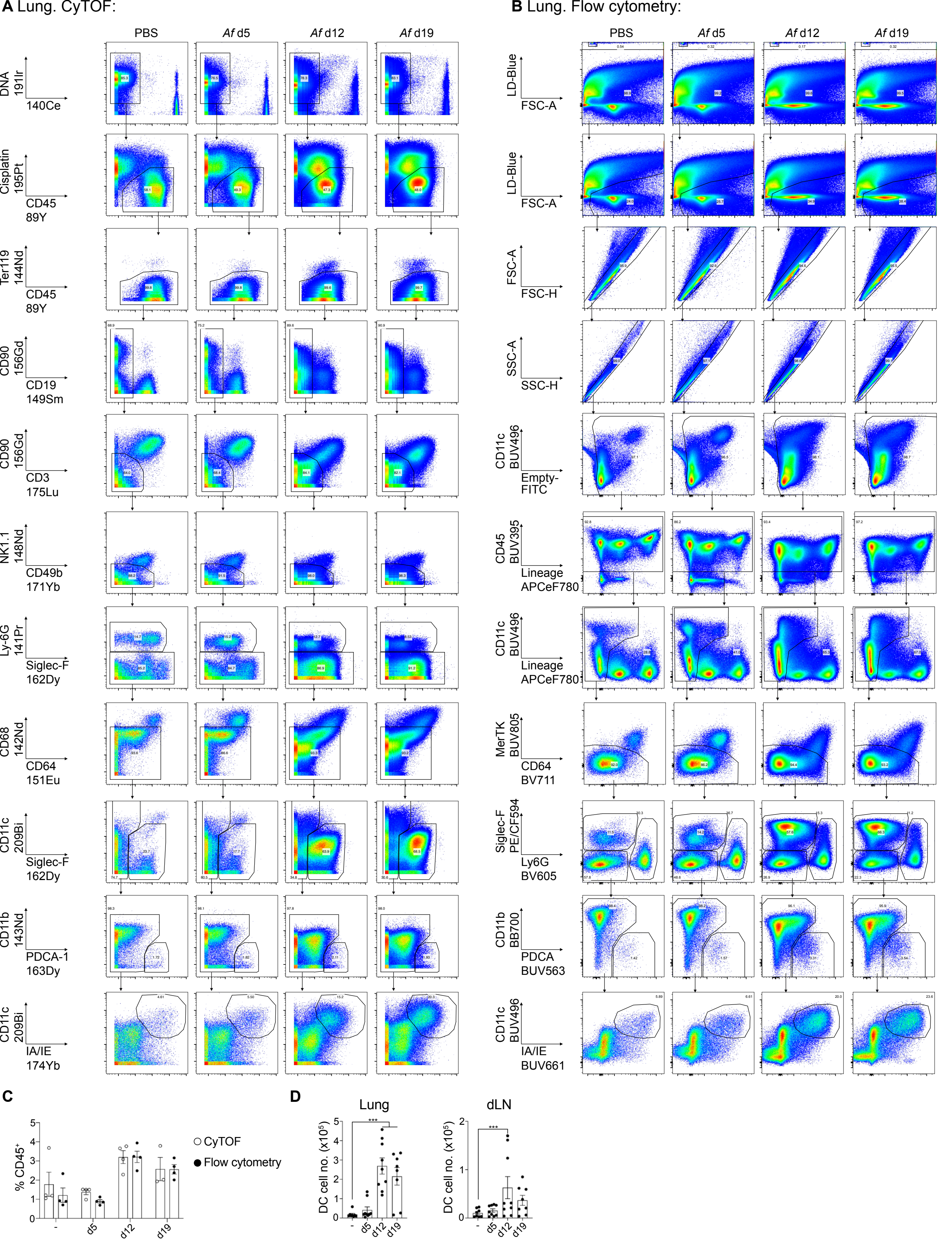


**Supplementary Figure 11**

### Supplementary Figure 12

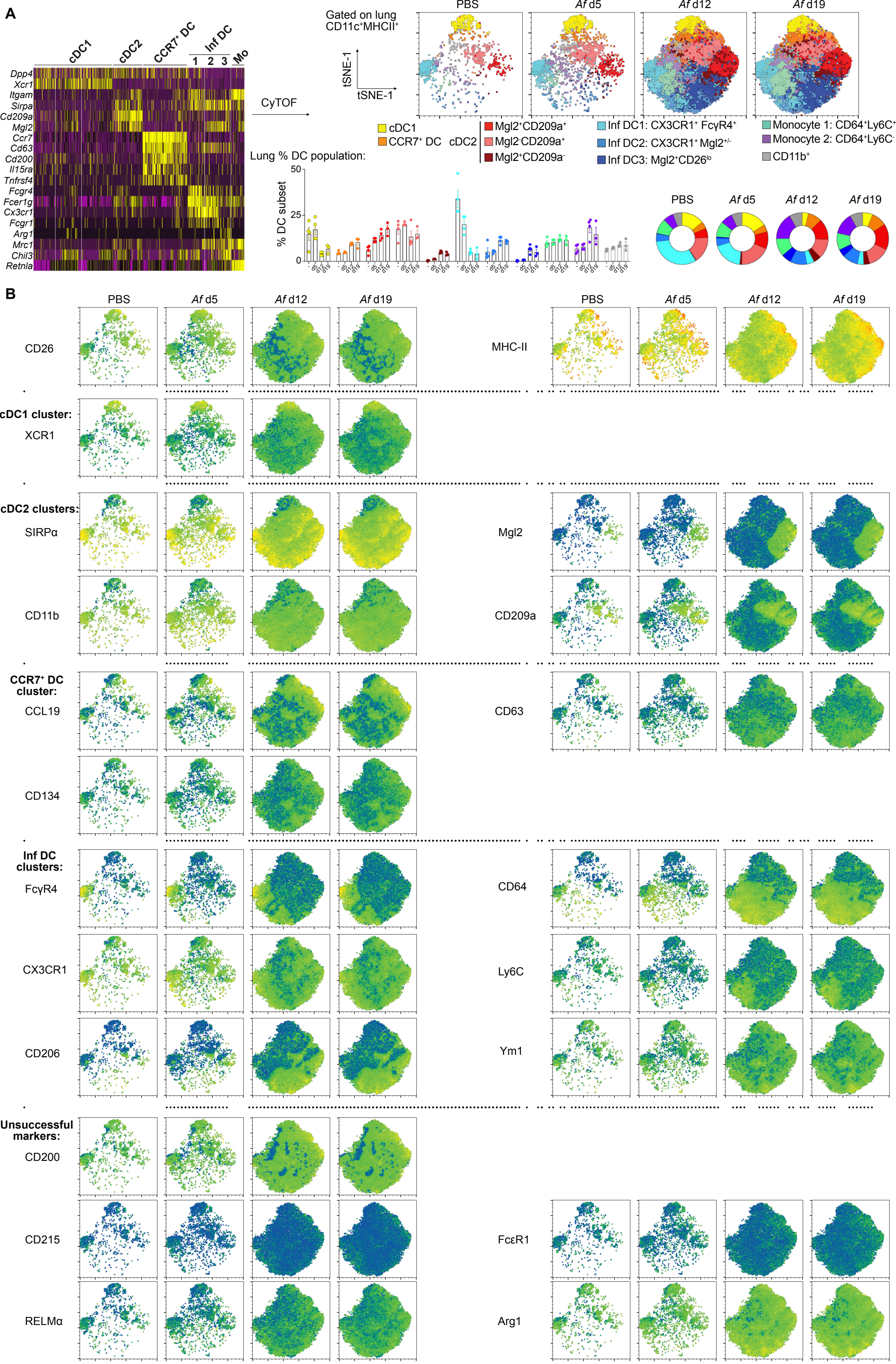


Supplementary Figure 12

### Supplementary Figure 13

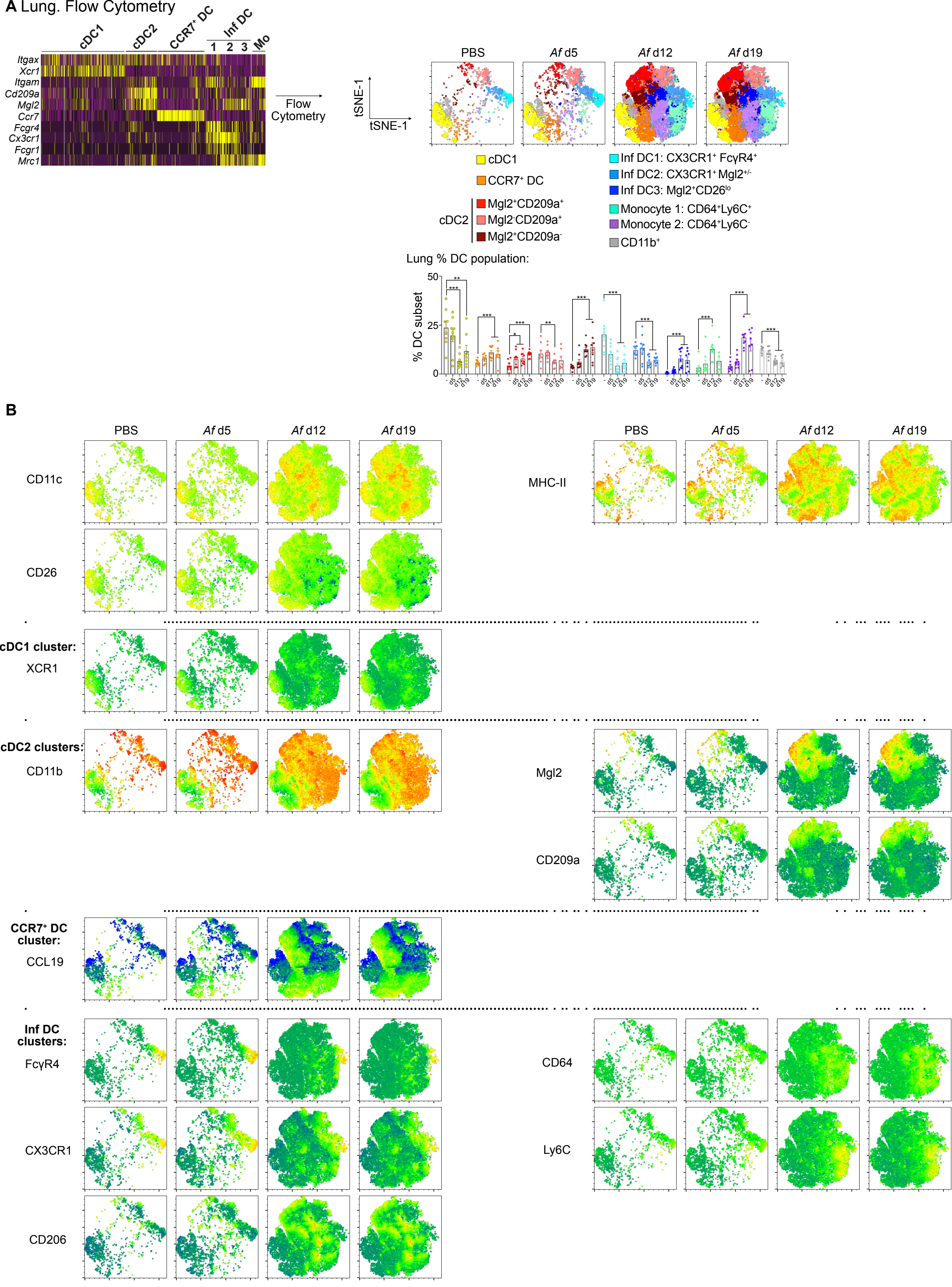


**Supplementary Figure 13**

### Supplementary Figure 14

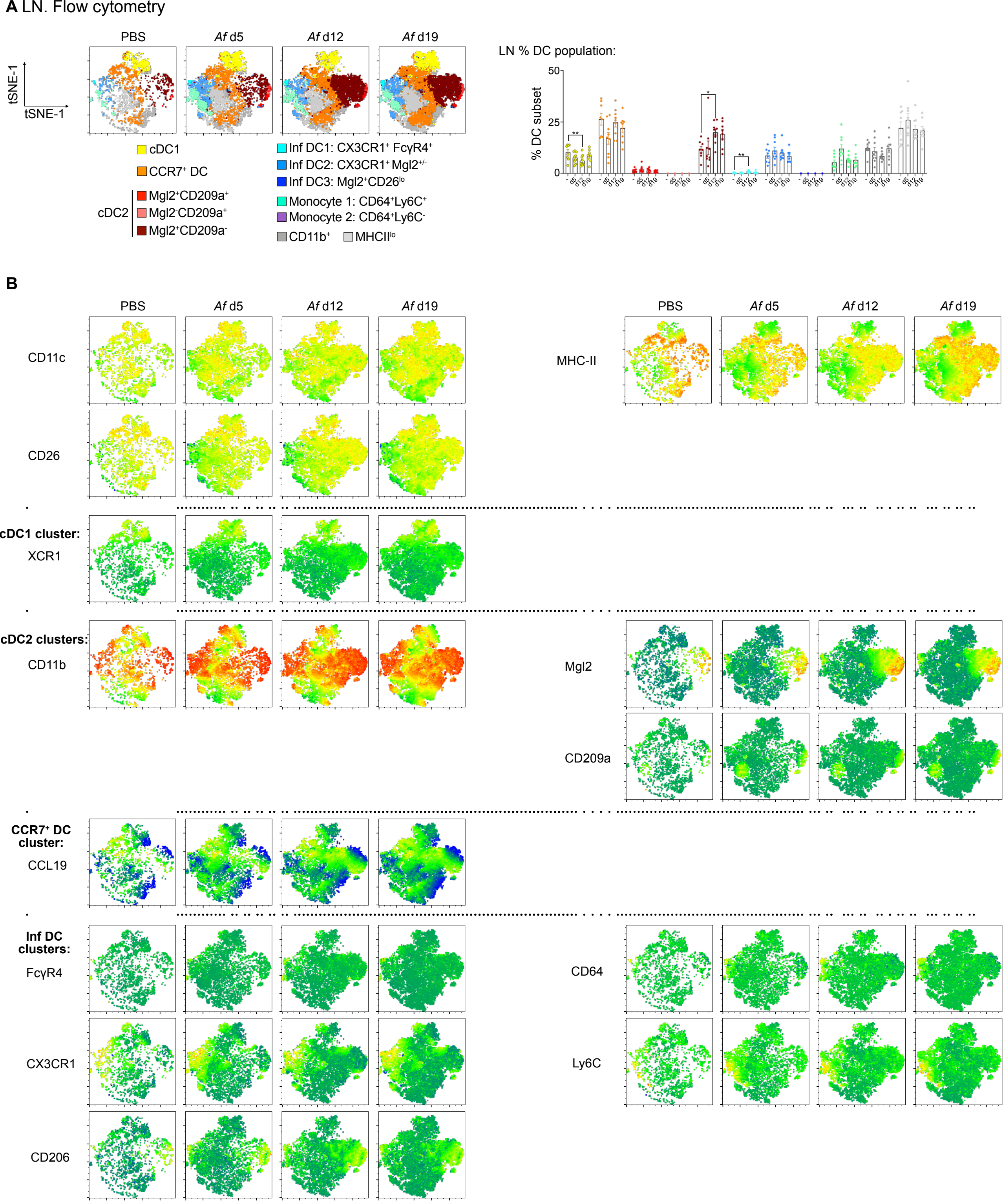


Supplementary Figure 14

### Supplementary Figure 15

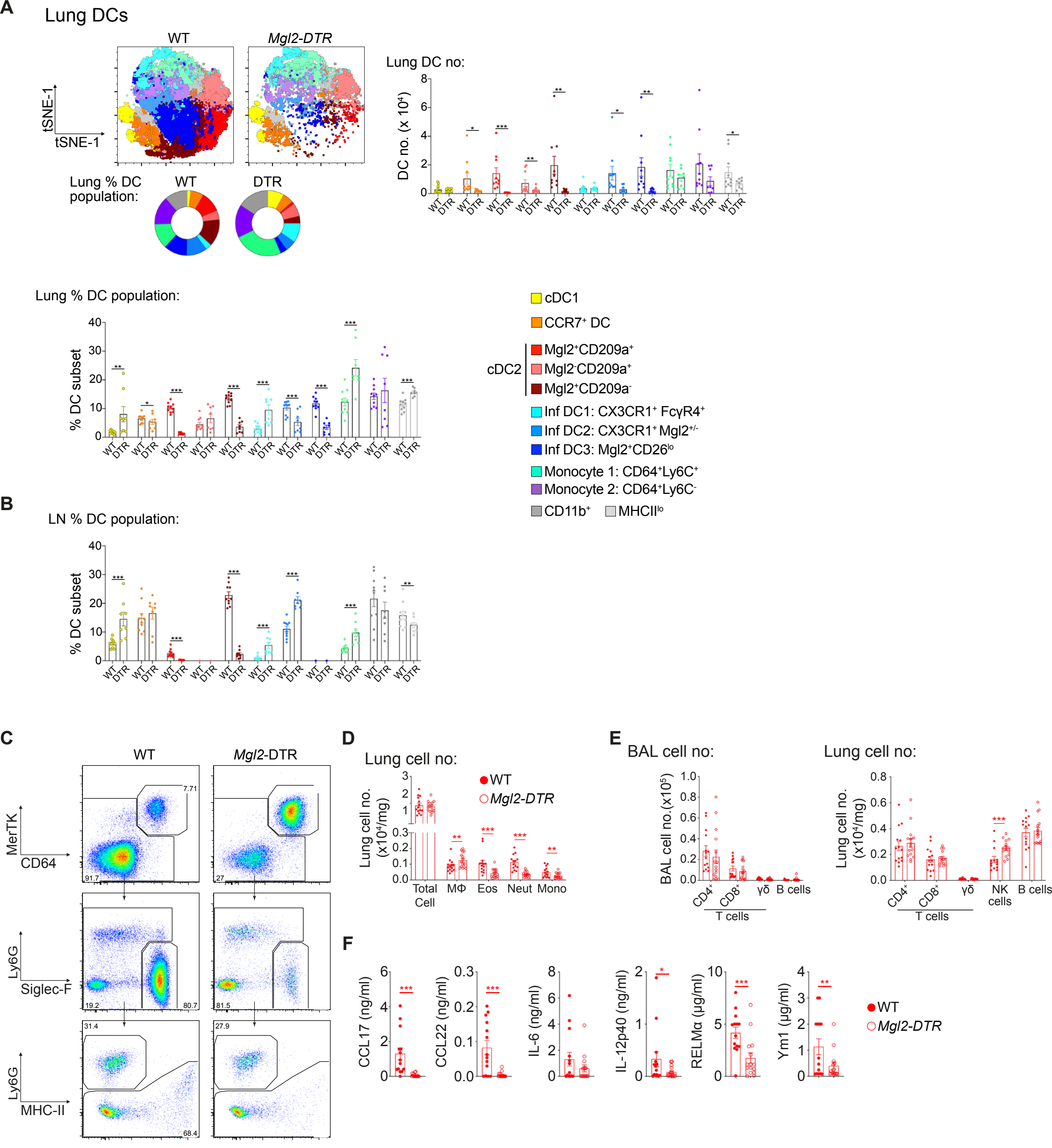


**Supplementary Figure 15**

### Supplementary Figure 16

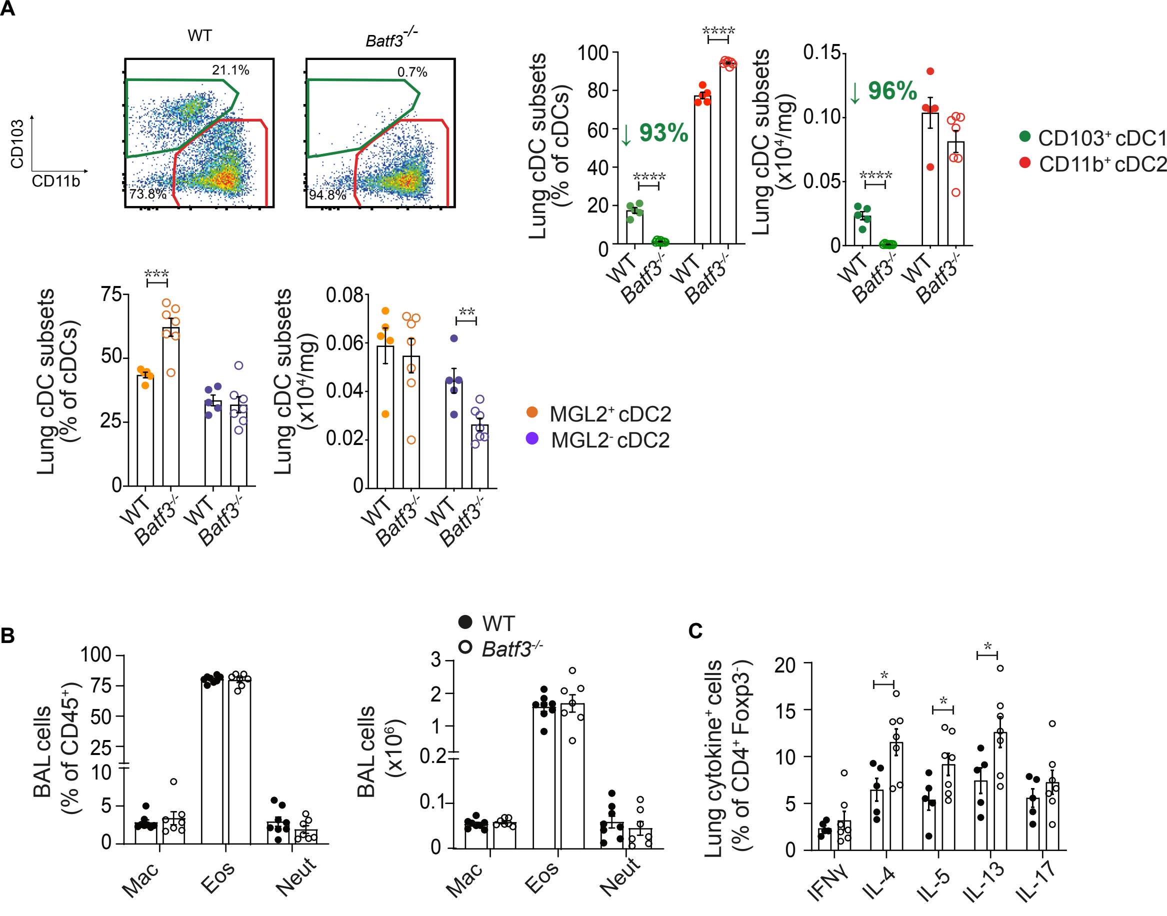


**Supplementary Figure 16.**
