## Supplementary Table 1 and 2 for "Mgl2^+^ cDC2s coordinate fungal allergic airway type 2, but not type 17, inflammation"

**Supplementary Table 1. List of qPCR primers**

| **Gene** | **Fwd** | **Rev** |
| --- | --- | --- |
| *Il4* | GAGAGATCATCGGCATTTTGA | TCTGTGGTGTTCTTCGTTGC |
| *Il5* | ACATTGACCGCCAAAAAGAG | CACCATGGAGCAGCTCAG |
| *Il10* | cagagccacatgctcctaga | Tgtccagctggtcctttgtt |
| *Il13* | CCTCTGACCCTTAAGGAGCTTAT | CGTTGCACAGGGGAGTCT |
| *Il17* | TGTGAAGGTCAACCTCAAAGTC | AGGGATATCTATCAGGGTCTTCATT |
| *Ifng* | GGAGGAACTGGCAAAAGGAT | TTCAAGACTTCAAAGAGTCTGAGG |

**Supplementary Table 2. List of flow and mass cytometry antibodies.**

| **Marker** | **Clone** | **Flow cytometry** | **Mass Cytometry** |
| --- | --- | --- | --- |
| Viability | - | Live Dead Blue | Cisplatin |
| CD45 | 30-F11 | BV510, BV785 or BUV395 | 89Y |
| CD3 | 17A2 | APC/eFluor780, Biotin | 175Lu |
| CD19 | 1D3 | APC/eFluor780, Biotin or PE/CF594 | 149Sm |
| CD90 | 30-H12 | APC/eFluor780 | 156Gd |
| NK1.1 | PK136 | APC/eFluor780, Biotin or PE/Cy5 | 148Nd |
| Ter-119 | TER119 | APC/eFluor780, Biotin | 144Nd |
| CD49b | DX-5 | - | 171A |
| CD45R (B220) | RA3-6B2 | Biotin |  |
| TCRb | H57-597 | APC/eFluor780 | - |
| TCRγδ | GL3 | PE | - |
| CD4 | RM4-5 | AF700 | - |
| CD8 | 53-6.7 | PE/Cy7 | - |
| IL-4 | 11B11 | PE/CF594 | - |
| IL-5 | TRFK5 | BV421 | - |
| IL-10 | JES5-16E3 | BV605 | - |
| IL-13 | eBio13A | AF488 | - |
| IL-17 | TC11-18H10.1 | PE/Cy7 | - |
| IFNγ | XMG1.2 | BV711 | - |
| F4/80 | BM8 | - | 155Gd |
| MerTK | 2B10C42 | APC, Biotin or FITC | anti-FITC 160Gd |
| CD64 | X54-5/7.1 | PE or BV711 | 151Eu |
| Siglec-F | E50-2440 | AF700, APC or PE/CF594 | anti-APC 162Dy |
| Ly-6G | 1A8 | APC/eFluor780, BV650, FITC or PerCP/Cy5.5 | 141Pr |
| Ly-6C | HK1.4 | AF700 or BV605 | 150Nd |
| CD11c | N418 | BV605 or BUV496 | 209Bi |
| IA/IE (MHC-II) | M5/114.15.2 | PE/Cy5 or BUV661 | 174Yb |
| PDCA-1 | 927 | BV650 or BUV563 | 163Dy |
| CD26 | H194-112 | BUV737 | 172Yb |
| CD11b | M1/70 | BB700 or BV711 | 143Nd |
| SIRPα | P84 | - | 173A |
| CD301b (MGL2) | URA-1 | PE or PE/Cy7 | anti-PE 165Ho |
| FcεR1 | MAR-1 | - | 176Yb |
| CD209a | MMD3 | PE | 152Sm |
| CD63 | NVG-2 | - | 146A |
| CD200 | OX2 | - | 158A |
| CD215 | 6B4C88 | - | 166A |
| CD134 | OX-80 | - | 168A |
| FcγR4 | 9E9 | BV421 | 154Sm |
| Arg1 | Polyclonal Sheep | - | 164Dy |
| RELMα | Polyclonal Rabbit | - | 159Tb |
| CX3CR1 | SA011F11 | PE/CF594 | 167Er |
| XCR1 | ZET | BV510 | 147Sm |
| CD103 | 2E7 | BV421 |  |
| CD206 | C068C2 | BV785 | 169Tm |
| Chi3l3/Ym1 | Polyclonal Goat | - | 161Dy |
| Slc7a2 | Polyclonal Rabbit | - | 153A |
